# Isoflurane and Injectable Anaesthesia Differentially Affect Optoretinography Signals in C57BL/6J Mice

**DOI:** 10.64898/2026.11.26.769405

**Authors:** Sybren Worm, Yuxiang Zhou, Georg Ladurner, Quang Nguyen, Theresa Rohrmoser, Lucas May, Maria Varaka, Yash Patel, Conrad Merkle, Katharina Heissl, Pengfei Zhang, Bernhard Baumann

## Abstract

**Purpose:** Anaesthesia is commonly required to immobilise mice in preclinical ophthalmic studies but also alters systemic physiology. We investigate the effects of three anaesthetics on optoretinography (ORG) measurements obtained by functional optical coherence tomography (OCT).

**Methods:** A total of 14 adult C57BL/6J mice (5 female and 9 male) were grouped in three cohorts. Each cohort was alternatingly exposed to isoflurane (ISO), ketamine-xylazine (KX) and a mixture of medetomidine, midazolam, fentanyl and ketamine (MMFK). ORG measurements were recorded by functional OCT 15 and 30 minutes after induction using a brief 502 nm light stimulus. The time to peak contraction, contraction amplitude and the elongation slope were extracted from ORG traces. Furthermore, a spectral analysis was performed on the respiratory rate acquired from the OCT data between both sexes. All parameters were statistically analysed using a linear mixed effects model with Holm-Bonferroni correction.

**Results:** Statistically significant differences were observed for some ORG parameters. The contraction amplitude for mice was greater and slope was lower under ISO anaesthesia compared to MMFK at 15 minutes post induction. Additionally, for both sexes, statistically significant differences were observed for the respiratory rate of mice anaesthetised with ISO compared to those under MMFK and KX.

**Conclusions:** All three anaesthetic protocols permitted ORG acquisition. However, anaesthesia-dependent effects on ORG parameters and respiratory rate should be considered when interpreting and comparing ORG data.

## Introduction

The photoreceptors in the retina are responsible for phototransduction, i.e. the transformation of photons into electrical signals, necessary for the image formation performed by the visual cortex in the brain. ^1^ In clinical ophthalmology, the function of the photoreceptors is monitored through a technique called electroretinography (ERG). ERG measures the electrical signals caused by phototransduction in response to light stimulation. ^2^ The subject is dark adapted and is then exposed to a light stimulus to induce the phototransduction. The resulting electrical signals are measured using an electrode, commonly placed close to the cornea. ^3^

In recent years, an optical analogue of ERG has been developed, namely optoretinography (ORG). ^4^ Instead of relying on electrodes, ORG optically assesses the function of the photoreceptors by measuring a change of the optical path length within the photoreceptors after a stimulus. ^5^ ORG is based on optical coherence tomography (OCT), a high-resolution imaging modality capable of recording volumes with micrometer scale resolution in real-time. In an ORG setup, an OCT system is combined with an extra light source that is used to stimulate the photoreceptors in sync with the data acquisition.^6^ The recorded optoretinogram depends on the excitation power, excitation wavelength and the pulse duration of the light flash inducing the phototransduction. ^7^ A typical ORG response of rods in rodents consists of an initial contraction (decrease of optical path length [OPL]) of the photoreceptors followed by an elongation of the photoreceptors. ^8^

Preclinical studies in mice are frequently performed in ophthalmic research to investigate specific elements of eye pathologies and gain an accelerated understanding about disease pathology. ^9^ Except for the clear difference in size and the lack of a macula, the mouse eye greatly resembles the human eye. Another major difference is the altered ratio between rods and cones in mice and humans. In mice, 97% of the receptors are rods, whereas the rod concentration in the human eye amounts to about 95%. ^10^ As the mouse retina is rod dominated, ORG response is optimized by choosing an excitation wavelength close to the sensitivity maximum of the rods close to 500 nm wavelength. ^11^

A common downside of ophthalmic research in mice is the need to immobilise mice with an anaesthetic agent in order to obtain stable images. While the relative effects on measurements between different anaesthesia methods remain largely unstudied, there have been some efforts to systematically test them. Recently, we investigated the effects of three different anaesthetic agents – namely isoflurane (ISO), ketamine-xylazine (KX) and a mixture of medetomidine, midazolam, fentanyl and ketamine (MMFK) – and their effects on the physiology and retinal dynamics in mice. ^12^ We observed that the choice of anaesthesia greatly affects the mouse physiology, such as the heart rate, blood pressure and blood vessel size. Nair et al. in 2011 reported a similar effect for the functional response of photoreceptors. Testing seven different anaesthetic protocols, including ISO and KX revealed a less pronounced ERG response in mice anaesthetised with ISO compared to those anaesthetised with KX. ^13^ Consistent with this, Ma et al. recently reported that anaesthesia alters mouse retinal structure and intensity-based ORG responses relative to awake imaging, with the largest differences between isoflurane and tribromoethanol and only small differences between isoflurane and esketamine+xylazine.^14^ However, the effects of ISO, KX, and MMFK on phase-based ORG readouts have not been directly compared.

Based on these findings, we hypothesise that ORG curves would similarly be reduced for ISO compared to MMFK and KX in the mouse eye. Therefore, in this work, we systematically investigate the effect of ISO, MMFK and KX on ORG measurements. We analyse the contraction amplitude, the time to peak contraction and the ORG slope after contraction both female and male C57BL/6J mice.

## Methods

### Mice and Experimental Procedure

A total of 14 C57BL/6J mice (The Jackson Laboratory, strain #000664) were randomly assigned to three groups, as shown in Fig. 1A. Group I consisted out of 3 males and 2 females, while groups II and III comprised 4 males and 1 female and 2 males and 2 females, respectively. For every group, a different anaesthetic agent was utilised for each of three weekly imaging sessions. Group I was anaesthetised with ISO for the first, KX for the second, and MMFK for the third imaging session. Group II and Group III followed a similar sequence, however, the initial imaging session was performed with KX and MMFK, respectively.

**Figure 1.**
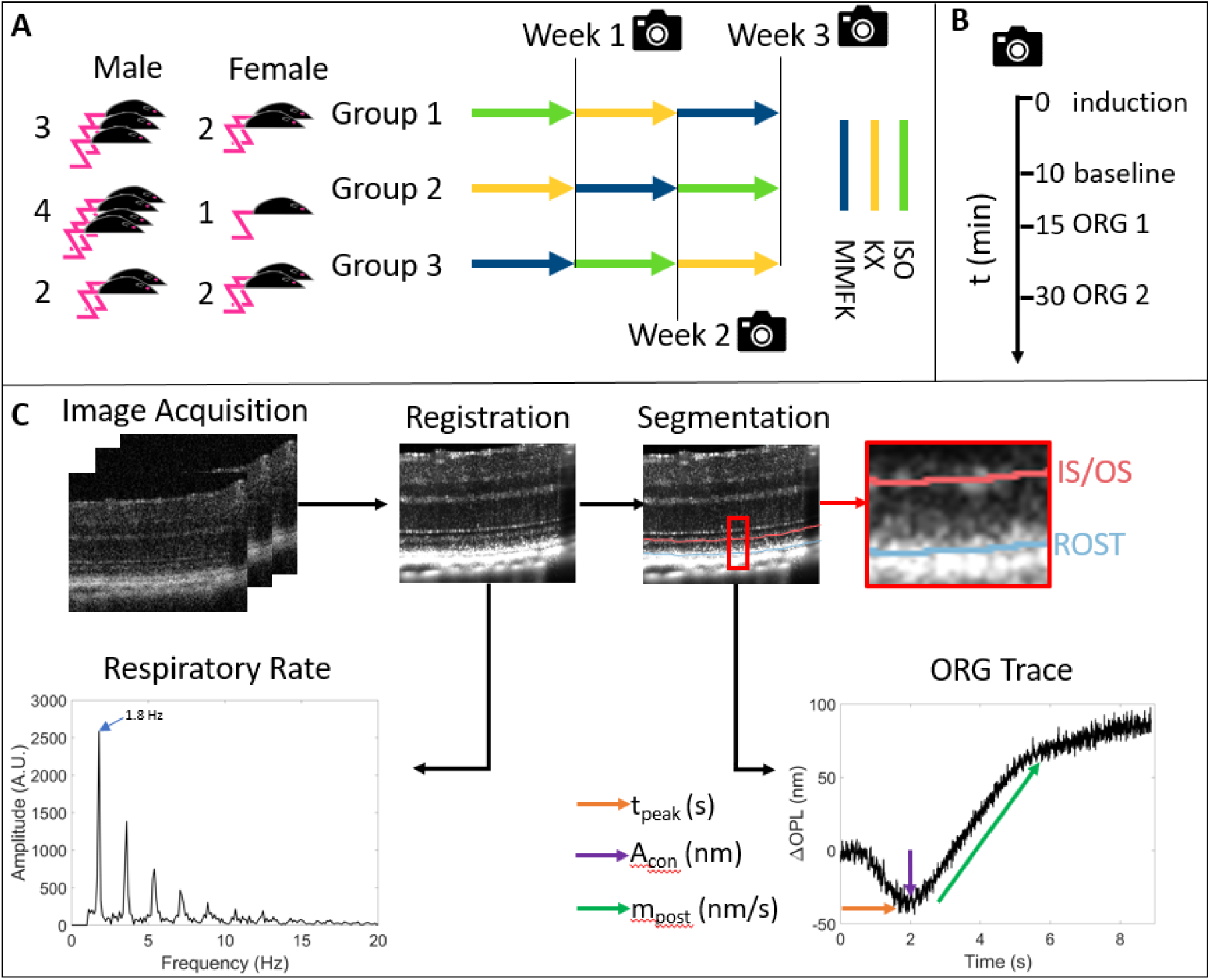
Experimental design and data processing workflow. A) Imaging protocol of the 3 different groups, depicting the order of anaesthetics the groups were exposed to and the number of mice per group. B) Timeline for each imaging session. C) Visual representation of the data processing steps.

Prior to each imaging session, the mice were dark adapted for a minimum of 2 hours. ^15^ The room was lit by LEDs emitting red light at 638-nm wavelength outside of the spectrum visually perceivable to mice, and any sources of light contamination were minimised to avoid breaking the dark adaptation. Afterwards, the mice were imaged individually. Each imaging session lasted 30 minutes, where t = 0 min denotes the moment a mouse was first exposed to an anaesthetic agent.

For imaging sessions with KX, the mice were injected intraperitoneally with a solution containing ketamine (100 mg/kg body weight) and xylazine (6.0 mg/kg). Similarly, MMFK was also administered as a solution containing medetomidine (0.3 mg/kg) midazolam (1.0 mg/kg), fentanyl (0.03 mg/kg) and ketamine (10.0 mg/kg) through an intraperitoneal injection. For both, KX and MMFK sessions, the mice were first placed in an induction chamber where the body temperature was maintained using an electric blanket. In the case for ISO, the mice stayed in the induction chamber for 4 minutes and were exposed to 4% ISO in oxygen. Once the mice lost the righting reflex and showed no visually observable ocular motion, they were transferred to the OCT imaging stage. There, the body temperature was maintained using iron powder-based hand warmers. In the case of ISO, the anaesthetic depth was maintained using 2% ISO in oxygen until the end of the imaging session. The right pupil of the mice was dilated using tropicamide eye drops (0.5%, Agepha Pharma s.r.o., Senec, Slovakia). During all imaging sessions, hydration eye drops (Théa Pharma GmbH, Berlin, Germany) were applied to the eyes of the mice. Any residual fluid was gently removed using a cotton swab just before an OCT dataset was acquired. Over the course of one imaging session, three OCT datasets were acquired. The first dataset, referred to as a baseline measurement, was taken around 10 minutes post anaesthesia induction and without an optical stimulation. The second OCT dataset was taken at 15 minutes post anaesthesia induction, including a light flash to stimulate the photoreceptors. Similarly, at 30 minutes anaesthesia post induction, a third OCT recording was taken to measure the recovery of the photoreceptors during the 15 minutes in between measurements. After the imaging session, the mice anaesthetised with MMFK received a partial antagonization consisting of atipamezole (1.0 mg/kg body weight) and flumazenil (0.1 mg/kg) injected intraperitoneal. On top of that the mice were also injected via a subcutaneous injection with 1 mL of a glucose solution to ease recovery. All procedures were approved by the local animal ethics committee and the Austrian authorities (protocol GZ 2024-0.044.300). All animal experiments were conducted in accordance with the ARVO Statement for the Use of Animals in Ophthalmic and Vision Research.

### OCT Ophthalmoscope and ORG Protocol

The data acquisition was performed with a custom-built polarisation sensitive OCT (PS-OCT) system as described by Fialová et al. ^16^ and modified as described by Harper et al. ^17^ The system operated at a central wavelength of 840 nm and a spectral bandwidth of 100 nm. The corresponding axial resolution was 5.1 μm in air or 3.8 μm in tissue. The device was operated at an A-scan rate of 80 kHz. The acquired datasets consisted of 2000 B-scans repeated in the same retinal location, and each B-scan of 256 A-scans. During alignment, the ONH was first centred in the 1 mm × 1 mm area provided by the real-time fundus preview. Afterwards, a B-scan was chosen up to 250 μm inferior to the ONH where the slow axis scanner was set stationary to scan a repeated B-scan pattern with a width of 0.5 mm. For the acquired datasets, the slow axis scanner was set stationary such that the same line was imaged over the course of 9 seconds. In addition, a custom LED ring consisting out of 12 LEDs in a half circle with a radius of 2 cm was placed on the sample arm objective situated about 2 cm away from the murine retina. A flash power of 26.6 μW was measured using a 2-mm diameter pupil at the position of the cornea. ^18^ In the non-baseline imaging sessions, the LED ring was used to provide an 0.33-s long optical stimulus at a wavelength of 502 nm triggered 1 second after the image acquisition had started.

### Image Processing

The raw spectral data obtained by the OCT system were processed as described in our previous work to obtain complex-valued OCT image data. ^19^ An analysis was performed to extract the respiratory rate based on our standard motion compensation algorithm. The algorithm smoothed the cross-polarised intensity data and binarized it using Otsu’s method. ^20^ Next an intensity weighted average was used to find the centroid of the data in the largest contiguous portion of the binarized mask, corresponding to the retinal pigment epithelium (RPE) and choroid. Finally, small scale fluctuations in the resulting centroid map were smoothed with a moving average. A Fourier analysis on the relative distance from the minimum centroid over all scans provided the respiratory motion, as illustrated in the example shown in Fig. 1B.

The ORG signals were extracted using a phase based method, as described in our previous work. ^21^ In short, subpixel motion correction was applied to both the amplitude and the phase components of the complex-valued OCT data. ^22^ The outer retinal layers, specifically the inner/outer segment junction (IS/OS) and the rod outer segment tips (ROST), were segmented on an averaged B-scan of the full dataset using a graph-based method, combined with manual corrections if needed. ^23^ The cross-spectrum between the two selected layers was then computed, and the corresponding time-dependent phase trace was extracted using a modified Knox-Thompson approach ^24^ to avoid phase wrapping and improve the signal-to-noise ratio (SNR), followed by spatial averaging and coherence-weight-based rejection and refinement of noisy points. Finally, the phase difference Δφ between the IS/OS and ROST was converted into layer displacement relative to t_0_ (t = 0 s) defined as the beginning of an acquisition of an OCT dataset using the following formula:

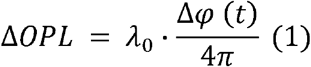

where λ_0_ is the central wavelength. After the ORG traces were extracted, the traces were smoothed using a moving average filter with a width of 50 pixels to reduce noise and the intercept at t_0_ was subtracted to start at ΔOPL(t_0_) = 0 nm. To quantify the ORG traces, the following three parameters were assessed. The contraction amplitude A_con_ was measured as the minimum ΔOPL of the ORG trace, whereas the time to peak contraction t_peak_ was defined as the corresponding time value, as shown in Figure 1C. Finally, the elongation slope m_post_ was determined by fitting a linear function from 0.44 seconds after t_peak_ for a total duration of 3.1 seconds. A spectral analysis was performed to identify frequency components within the baseline measurements. The baseline traces were Fourier transformed and the median noise floor for all anaesthetics was determined. Then the area under the curve (AUC) was determined as a summation of all areas above and below the curve with respect to the noise floor in the frequency range of roughly one standard deviation of the respiratory rate centred at the peak detection frequency.

### Statistical Analysis

To test for statistical differences, linear mixed effects models were applied to the measurements of the contraction amplitude, contraction time and the slope of the ORG recordings as well as the respiratory rate and the AUC for the respiratory rate and the heart rate. The majority of measurements were pooled together, whereas the respiratory rate was analysed for all anaesthetics and sex separately. The variable effect for this analysis was the respective anaesthesia. Where applicable, repeated measurements were taken into account by modelling a random intercept for each mouse. The maximum likelihood method was used to estimate the models and the method successfully converged for all models. The level of significance was set at a value of p = 0.05.

The analysis was separated into two different families: (i) the primary research parameters and (ii) the secondary investigated parameters. The primary research question entails the ORG parameters, i.e. the time to peak contraction (t_peak_) the contraction amplitude (A_con_) and the elongation slope post contraction (m_post_). The remaining parameters were part of the secondary investigated parameters, such as the respiratory rate, the AUC and the changes between both ORG recordings. Each family had an independent Holm-Bonferroni post correction applied to them to adjust for the multiple tests. The ORG parameters were statistically tested for the different anaesthetics at 15 minutes after anaesthesia induction. The measurements at 30 minutes after anaesthetic induction were used to test for differences compared to their counterparts assessed at 15 minutes for the same anaesthesia.

## Results

### Optoretinography

ORG traces were acquired in the right eyes of 14 C57BL/6J mice. Figure 2A shows the individual smoothed baseline measurements performed in order to determine the expected noise levels, while Fig. 2B shows the averaged traces for each anaesthetic. The green, blue and yellow lines represent the measurements obtained for the individual eyes with ISO, MMFK and KX, respectively, and the shaded areas represent the standard deviation. In this plot, the ORG data, from 47 male datasets and only 18 female datasets obtained from the 14 eyes are displayed. The low number of female datasets was due to the relatively high failure rate with female mice, as shown in Supplementary Table 1. Reasons for dropout were artifacts caused by strong eye motion, poor image quality, and vasculature masking a large portion within the B-scan. Figure 2C depicts smoothed ORG measurements acquired at 15 minutes after induction of the anaesthetic agent, whereas Fig. 2D shows the averaged traces. The 0.33-second long light stimulus fired 1 second into the recording is visualized by the vertical green and red line marks. Similarly, Figs. 2E and 2F show the individual smoothed traces and the average traces at 30 minutes, respectively.

**Figure 2.**
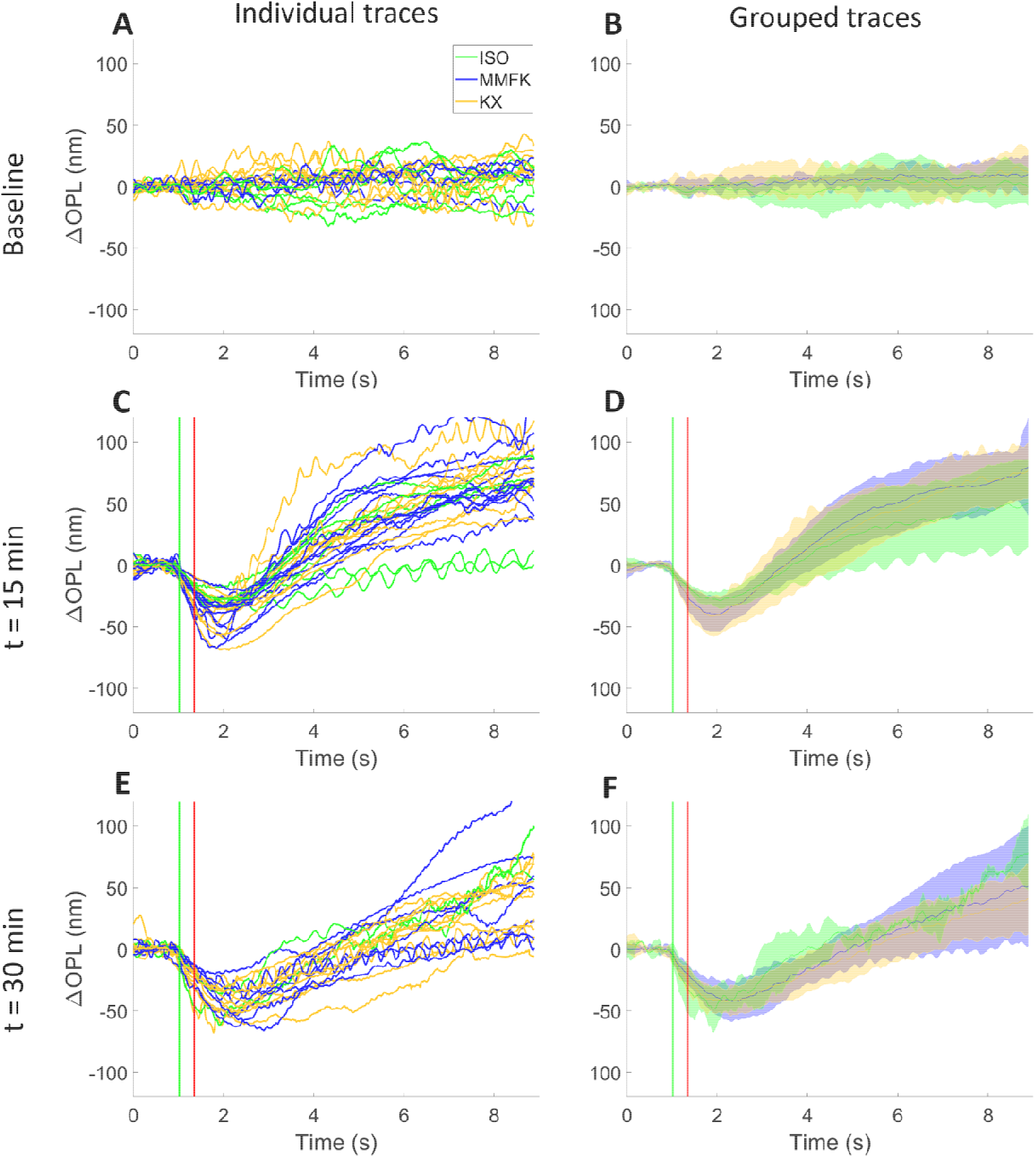
ORG for ISO (green), MMFK (blue) and KX (yellow). The green line at t = 1 s and the red line at t = 1.33 s mark the start and the end of the excitation flash, respectively. A) Baseline measurements without any stimulus. B) Averaged ORG traces from panel A, where the shaded areas indicate the standard deviation (STD) around the average curves (mean ± STD). C) ORG responses at 15 minutes post anaesthesia induction. D) Averaged ORG traces at t = 15 min. E) ORG traces at 30 minutes post anaesthesia induction, i.e. with 15 minutes recovery after the first flash. F) Averaged ORG traces at t = 30 min.

Three different attributes of the ORG curves – namely, t_peak_, A_con_, and m_post_ – were quantitatively assessed and plotted in Fig. 3. t_peak_ is shown in Figs. 3A and 3B at 15 minutes and 30 minutes, respectively. Figures 3C and 3D show A_con_ for the ORG measurements at 15 minutes and 30 minutes, respectively. Similarly, Figs. 3E and 3F show m_post_ for both stimulus experiments, respectively. Supplementary Table 2 provides the results of all statistical tests run to analyse differences between the various quantities in more detail. Of note, for t_peak_, no statistically significant difference was observed (p > 0.05). However, the ORG slope acquired at t = 15 min was statistically significant different between ISO and MMFK (p = 0.02). Also, for A_con_, a significant difference was observed comparing ISO and MMFK (p = 0.006) whereas no statistically significant differences were found when comparing KX to ISO and MMFK to KX, respectively (p > 0.05).

**Figure 3.**
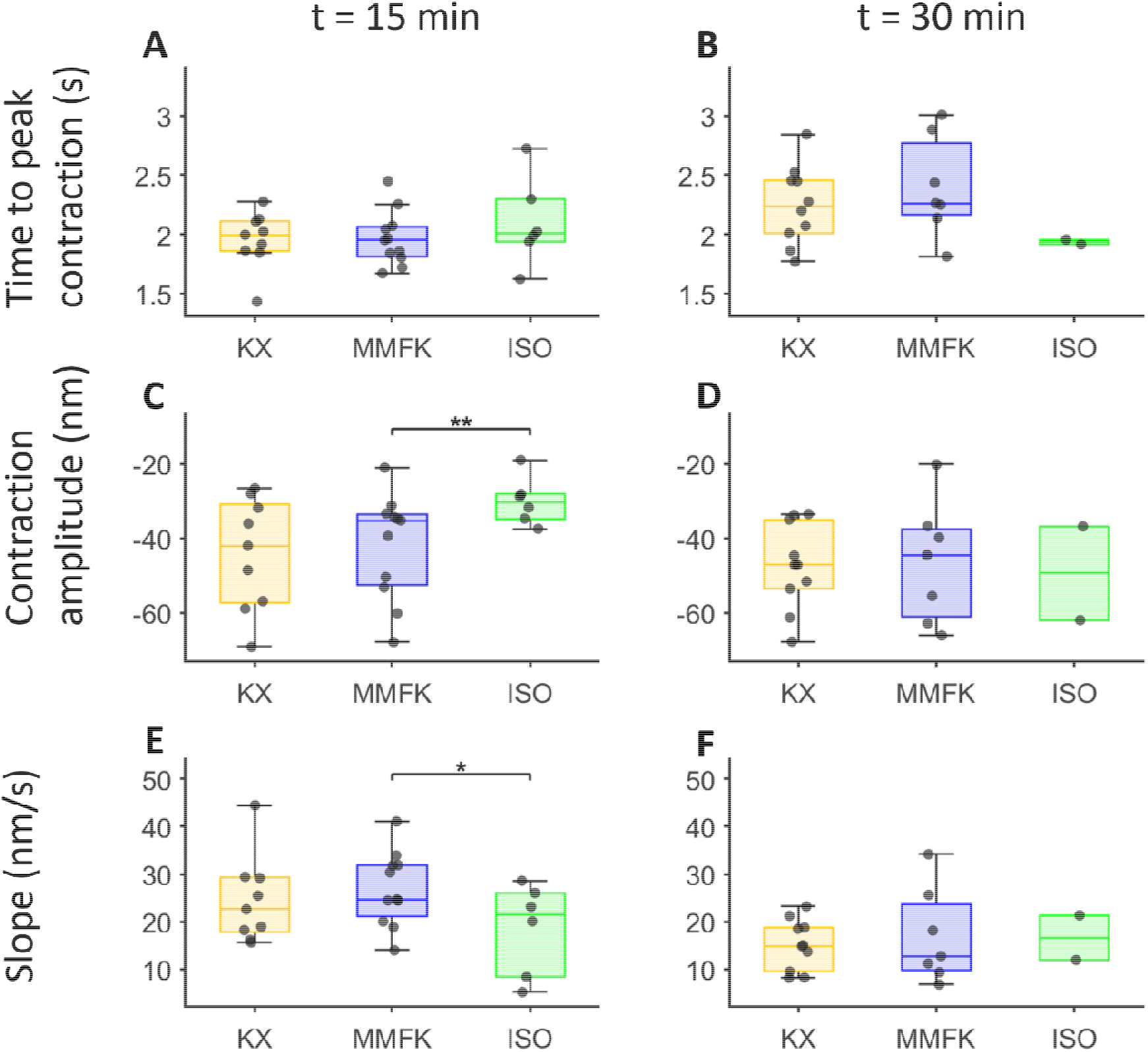
Visualisation of the statistical analysis in box plots, where yellow, blue and green correspond to KX, MMFK and ISO, respectively. A, B) Time to peak contraction for the ORG measurements at t = 15 min and t = 30 min, respectively. C, D) Contraction amplitude for all anaesthetics for the ORG measurement at t = 15 min and t = 30 min, respectively. E, F) The corresponding slope after the contraction for these ORG measurements at t = 15 min and t = 30 min, respectively. Asterisks indicate significant differences between MMFK and ISO after Holm–Bonferroni correction (*p < 0.05, **p < 0.01).

Generally, lower slopes can be observed for the set of ORG traces obtained after the second stimulus (t = 30 min) compared to the first stimulus (t = 15 min) as shown in Figs. 3E and 3F. Within the MMFK group, the slope differed significantly between the 15-min and 30-min recordings (p = 0.02), whereas no significant within-group difference was found for ISO (p = 0.75) or KX (p = 1).

### Respiratory Rate

Figure 4A shows an example baseline measurement of a mouse anaesthetised with MMFK. In Fig. 4B, the Fourier spectrum of the baseline measurement shown in Fig. 4A visualises the contribution of frequency components. The blue shaded area is the AUC of the respiratory rate for a duration corresponding to the standard deviation of the respiratory rates among all measurements per anaesthetic as determined from Fig. 4C. Figure 4C visualises the extracted respiratory rates for each sex and anaesthetic in boxplots. The mean respiratory rate for females was 173.9 ± 26.0 breaths per minute (BPM), 175.0 ± 22.8 BPM and 130.5 ± 24.5 BPM for KX, MMFK and ISO, respectively. The mean respiratory rate for the male mice was 195.2 ± 21.8 BPM, 191.3 ± 16.4 BPM and 138.9 ± 22.6 BPM. For both male and female mice, a significantly lower respiratory rate was found for ISO compared to MMFK and KX (p < 0.0001). In Fig. 4D, the average frequency response per anaesthetic is shown. AUC of the respiratory peak did not differ significantly among anaesthetics (all adjusted p = 1); Averaged AUC values were 58% and 34% higher for KX and MMFK than for ISO, respectively.

**Figure 4.**
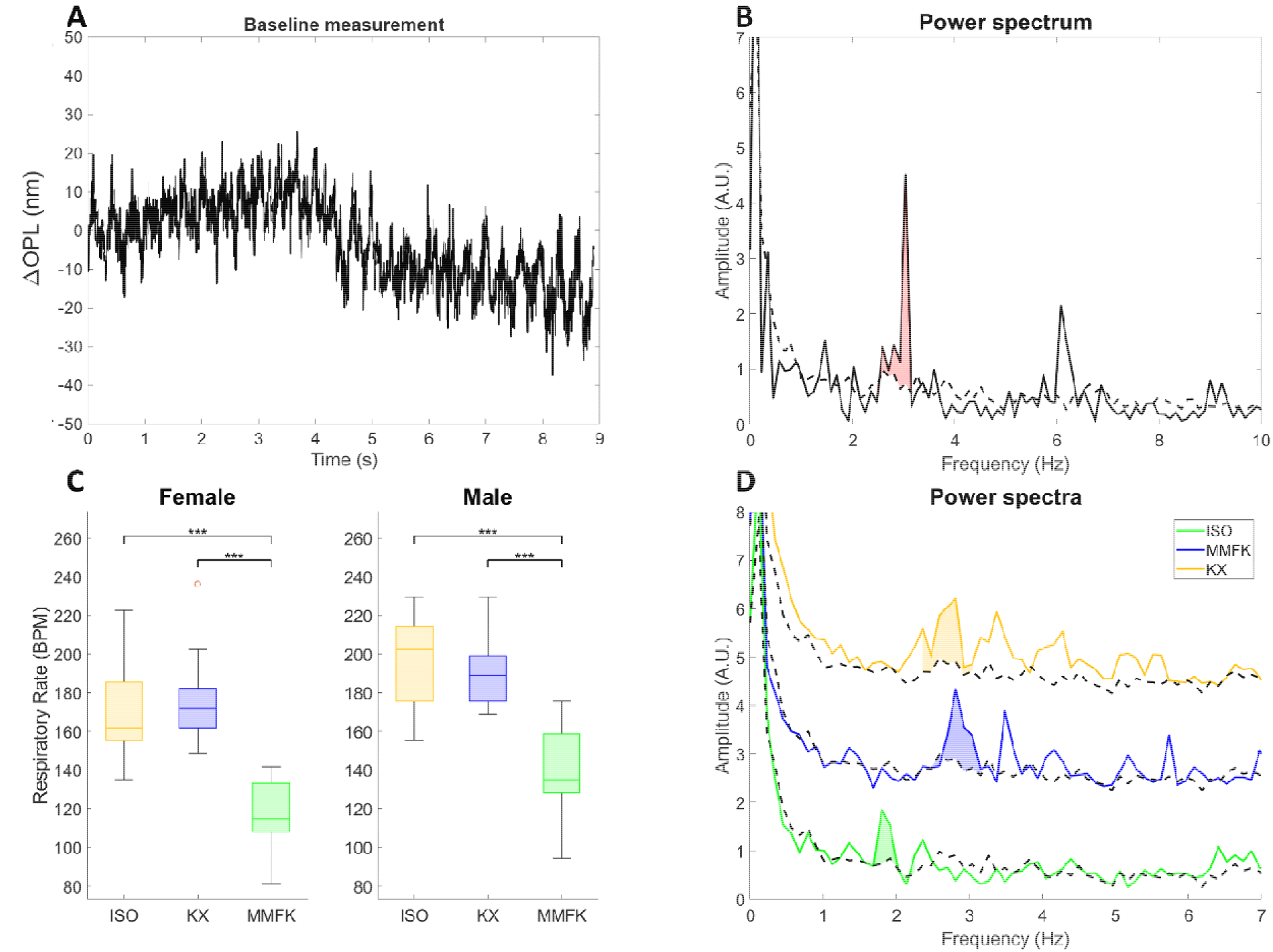
Analysis of breathing related motion patterns in ORG traces. A) Example baseline measurement. B) Power spectrum of the baseline measurement in panel (A) where the red area indicates the expected range covered by the respiratory rate and the dashed line is the median power spectrum over all anaesthetics. C) Respiratory rate for the three anaesthetics stratified into female and male mice. D) Averaged power spectra for the three anaesthetics. The shaded areas indicate the average AUC for each anaesthetic. The spectra for MMFK and KX were plotted with vertical offsets of 2 and 4 A.U. to enhance the visibility. The dashed line indicates the frequency content of the background and was computed as the median power spectrum taken over all anaesthetics.

## Discussion

This work provided a direct comparison between three anaesthetic agents and explored the influence of different anaesthesia on ORG measurements and respiratory rate. 4 out of 6 the investigated parameters extracted from the ORG traces did not reveal any statistically significant differences (cf. Figs. 2 and 3) and thus indicated a comparable ORG performance for all three anaesthetics. However, the maximum contraction amplitude was statistically significantly different for ISO compared to MMFK. The smaller peak amplitude for ISO might be related to a lower photoisomerization, i.e. an impeding effect of ISO on the function of the photoreceptors. ^25^ Such an impeding effect of ISO on phototransduction would also support our initial hypothesis and thereby be in tandem with the work by Nair and co-workers who observed a lessened ERG response for ISO compared to KX. ^13^ In our study, KX did not differ significantly from the other anaesthetics regarding the contraction amplitude, albeit a larger sample size or a more lenient statistical post correction for the studies might have changed this outcome.

Our finding of an ISO–MMFK difference is broadly consistent with Ma et al., who observed anaesthesia-dependent changes in mouse ORG signals. However, they found only small differences between ISO and esketamine+xylazine, whereas we did not observe a significant ISO–KX difference for most ORG parameters. These discrepancies may reflect differences in anaesthetic agent (esketamine vs. ketamine), ORG contrast (intensity-based vs. phase-based), stimulus parameters, time after induction, and sample size^14^

Furthermore, Figs. 2C and 2D revealed a difference between ORG at 15 min post induction with over 2 hours of dark adaptations compared to the subsequent ORG measurement taken an additional 15 minutes later. One possible interpretation is a slower post-stimulation recovery under MMFK, although the present data cannot distinguish this from anaesthesia duration, drift, or sample-related effects. While all of the three anaesthetics used in this work proved a suitable choice to perform ORG measurements with, the eventual dependence of the ORG response on anaesthesia protocol and time after induction needs to be considered carefully when designing experiments and comparing ORG data obtained with different methods.

For the measurements where the mice were anaesthetised with ISO, a higher dropout rate compared to MMFK and KX was observed (cf. Supplementary Table 1). The main reason for dropout in our experiment was strong eye motion under ISO which defeated the ORG analysis in 52% of the failed datasets. In a study performed in awake and anaesthetised mice using KX, Feng et al. reported that KX supressed fast eye motion. ^26^ Moreover mice anaesthetised with ISO were reported to have more eye drift compared to KX which could in part explain the relatively high dropout ratio we observed for ISO. ^13^ ISO is, however, considered a safer anaesthetic option compared to KX. ^27^ KX is known for having a relatively higher mortality rate, but also bringing a more consistent anaesthetic depth. ^27,28^ While under KX the physiological parameters did stay more stable over the time span of 40 minutes, drop out due to KX was also observed in our cohort, as can be seen in Supplementary Table 1. The use of MMFK as an alternative anaesthetic agent for KX that is partially antagonizable is recommended by the Medical University of Vienna (MUW). However, since to the best of our knowledge, the use of MMFK is limited to the MUW, only a sparse amount of comprehensive and comparative research data is currently available which limits the understanding of MMFK.

Another factor that warrants further investigation is the larger relative difference of unsuccessful measurements between male and female mice among all anaesthetics (cf. Supplementary Table 1). A plausible explanation may be an inherently lower anaesthetic sensitivity in female mice. ^29^ A lower anaesthetic depth with substantial eye motion would likely lead to more out of plane drift during repeated B-scan acquisitions and result in a degradation and even loss of the ORG signal. In future work, it could be worth monitoring the anaesthetic depth via additional parameters such as blood pressure to further explore the time characteristic of differences in the ORG response related to anaesthesia protocol and sex.

Statistically significant differences were also found for the respiratory rate for ISO compared to MMFK and KX for both sexes. These observations align with differences between respiratory parameters under ISO and KX reported in the literature, although the respiratory rates we measured were on average roughly 50 BPM higher. ^30^ Schmitz et al. assessed the respiratory rate for KX, ISO and a mixture of midazolam, medetomidine and fentanyl (MMF) in guinea pigs. Although direct comparisons cannot be made due to dissimilar species, different concentration of midazolam, medetomidine and fentanyl and the lack of ketamine in MMF, the authors reported a general trend similar to our findings with significantly lower respiratory rates for ISO compared to KX and MMF(K). ^31^

A spectral analysis of the ORG data also enabled the simultaneous measurement of the respiratory rate (cf. Fig. 4C). The AUC of the respiratory rate was shown to have large percentual differences between the anaesthetics, although not statistically significant. The increased AUCs for KX and MMFK indicate that the corresponding ORG measurements may be influenced more strongly by the respiratory motion. Chung et al. investigated the variability of the respiratory rate for both KX and ISO in rats and found that the variability remains similar for both anaesthetics. ^32^ One possible explanation for the lower AUC of the respiratory rate in the case for ISO compared to KX and MMFK might be that a lower frequency term might be better compensated by the registration software. Moult et al. performed a comparative study using rats and five anaesthetics, including KX and ISO. Part of the investigated criteria was the amount of drift in the retinal OCT data, which was higher for the rats anaesthetised with ISO compared to KX. Another investigated parameter was the time stability of the blood flow over a time period of 20 minutes, which was stable for ISO, whereas KX was not stable over this time period. ^33^ If the same holds true for mice, this could be an explanation why more measurements with ISO failed compared to KX but have less spread in the spectral analysis.

While this study revealed obvious differences of ORG parameters and respiratory rate between different anaesthetic protocols, its scope towards a sex-stratified analysis is limited owing to the relatively small sample size. Especially considering the larger failure rate observed in female mice. Similarly, the relatively high failure rate for ISO was not anticipated, in particular for the second ORG measurement. In future studies, an increased sample size could thus explore longitudinal differences of the optoretinogram related to the anaesthesia method as well as sex dependent differences of the functional response – an aspect that to the best of our knowledge has not been investigated using ORG thus far.

For ORG studies in mice, all three protocols proved feasible. However, ISO was associated with more motion-related failures and altered ORG amplitude, whereas KX and MMFK provided more stable recordings at the cost of greater respiratory motion and, for KX, higher safety concerns. In light of their eventual impact on the measured ORG response, anaesthetic protocol and time after induction should therefore be reported and need to be considered when comparing ORG data.

## Acknowledgements

The authors thank Sonja Reynoso-De-Leon, Christian Schönauer and the animal facility team for their technical support.

