## Supplements for "Isoflurane and Injectable Anaesthesia Differentially Affect Optoretinography Signals in C57BL/6J Mice"

**Supplementary Material**

**Supplementary Table S1.** An overview of the total datasets and the numbers of datasets that were unsuccessfully analysed using for each sex and the corresponding anaesthetic agent.

| **Anaesthetic**  **Sex** | **ISO**  (unsuccessful / total) | **MMFK**  (unsuccessful / total) | **KX**  (unsuccessful / total) |
| --- | --- | --- | --- |
| **Female** | 11/14 | 6/14 | 8/15 |
| **Male** | 10/21 | 5/21 | 5/26 |

**Supplementary Table S2.** An overview of the measured parameters, indicating the investigated parameter, the pair wise comparison made within the parameter, the family indicating the post hoc compensation and the adjusted p-value. Respiratory rate is abbreviated as RR.

| **Parameter** | **Comparison** | **Family** | **p-value (Holms)** |
| --- | --- | --- | --- |
| t_peak_ | ISO - KX | Primary | 0.54 |
| t_peak_ | ISO - MMFK | Primary | 0.25 |
| t_peak_ | KX - MMFK | Primary | 0.50 |
| A_con_ | ISO - KX | Primary | 0.20 |
| A_con_ | ISO - MMFK | Primary | 0.006 |
| A_con_ | KX - MMFK | Primary | 0.32 |
| m_post_ | ISO - KX | Primary | 0.50 |
| m_post_ | ISO - MMFK | Primary | 0.03 |
| m_post_ | KX - MMFK | Primary | 0.32 |
| AUC RR | ISO - KX | Exploratory | 1 |
| AUC RR | ISO - MMFK | Exploratory | 1 |
| AUC RR | KX - MMFK | Exploratory | 1 |
| RR Female | ISO - KX | Exploratory | 1.2×10^-11^ |
| RR Female | ISO - MMFK | Exploratory | 3.7×10^-12^ |
| RR Female | KX - MMFK | Exploratory | 1 |
| RR Male | ISO - KX | Exploratory | 1.7×10^-15^ |
| RR Male | ISO - MMFK | Exploratory | 5.9×10^-14^ |
| RR Male | KX - MMFK | Exploratory | 1 |
| RR | Female - Male | Exploratory | 1 |
| RR | Female – Male | Exploratory | 0.25 |
| RR | Female - Male | Exploratory | 0.45 |
| m_post_ | ISO ORG1 – ORG2 | Exploratory | 0.75 |
| m_post_ | KX ORG1 – ORG2 | Exploratory | 1 |
| m_post_ | MMFK ORG1 – ORG2 | Exploratory | 0.02 |
| A_con_ | ISO ORG1 – ORG2 | Exploratory | 1 |
| A_con_ | KX ORG1 – ORG2 | Exploratory | 1 |
| A_con_ | MMFK ORG1 – ORG2 | Exploratory | 1 |
| t_peak_ | ISO ORG1 – ORG2 | Exploratory | 0.41 |
| t_peak_ | KX ORG1 – ORG2 | Exploratory | 1 |
| t_peak_ | MMFK ORG1 – ORG2 | Exploratory | 1 |
